# Hypoxia-Sonic Hedgehog Axis as a Driver of Primitive Hematopoiesis Development and Evolution in Cavefish

**DOI:** 10.1101/2024.06.09.598120

**Authors:** Corine M. van der Weele, Katrina C. Hospes, Katherine E. Rowe, William R. Jeffery

**Affiliations:** Department of Biology, University of Maryland, College Park, MD 20742. USA

**Keywords:** *Astyanax mexicanus*, cavefish, primitive hematopoiesis, erythrocyte development, evolution, hypoxia, Sonic hedgehog

## Abstract

The teleost *Astyanax mexicanus* consists of surface dwelling (surface fish) and cave dwelling (cavefish) forms. Cavefish have evolved in subterranean habitats characterized by reduced oxygen levels (hypoxia) and show constructive and regressive phenotypic traits controlled by increased Sonic hedgehog (Shh) signaling along the embryonic midline. The enhancement of primitive hematopoietic domains, which are formed bilaterally in the anterior and posterior lateral plate mesoderm, are responsible for the development of more larval erythrocytes in cavefish relative to surface fish. In this study, we determine the role of hypoxia and Shh signaling in the development and evolution of primitive hematopoiesis in cavefish. We show that hypoxia treatment during embryogenesis increases primitive hematopoiesis and erythrocyte development in surface fish. We also demonstrate that upregulation of Shh midline signaling by treatment with the Smoothened agonist SAG increases primitive hematopoiesis and erythrocyte development in surface fish, whereas Shh downregulation via treatment with the Smoothened inhibitor cyclopamine decreases these traits in cavefish. Together these results suggest that hematopoietic enhancement is regulated by hypoxia and the Shh signaling system. Lastly, we demonstrate that hypoxia treatment enhances expression of Shh signaling along the midline of surface fish embryos. Thus, we conclude that a hypoxia-Shh axis may drive the adaptive evolution of primitive hematopoiesis and erythrocyte development in cavefish.

**Highlights:**

- Hypoxia increases hematopoiesis and erythrocytes in surface fish
- Shh upregulation increases hematopoiesis and erythrocytes in surface fish
- Shh inhibition decreases hematopoiesis and erythrocytes in cavefish
- Hypoxia upregulates Shh along the embryonic midline in surface fish

## Introduction

The teleost *Astyanax mexicanus* is an excellent model to study the evolution of development (Jeffery, 2001; 2020; Perera et al. 2023). A key feature of this model is the existence of different morphs of the same species, surface fish and cavefish, which recently diverged from a common ancestor (Fumey et al., 2018; Herman et al., 2018) and show multiple phenotypic differences. Over the past 20,000-300,000 years, cavefish have evolved regressive traits, such as eye degeneration, albinism or reduced pigmentation, and the loss of visually mediated behaviors, and constructive traits, such as the enhancement of non-visual sensory systems, oral development, olfaction, adipose deposition, and the appearance of novel feeding behaviors (Jeffery, 2020; Kowalko, 2020; Perera et al. 2023). The *Astyanax* system provides an opportunity to understand the genetic and molecular underpinnings of cavefish trait evolution. Some traits, such as the degeneration of eyes, are controlled by many genes, whereas others, such as albinism, are controlled by a single gene, and the underlying gene mutations can be in coding or non-coding regions (Protas et al., 2006; Hinaux et al., 2013; Ma et al., 2020). One of the most striking molecular differences between surface fish and cavefish is in the intensity of the Sonic Hedgehog (Shh) expression along the embryonic midline (Yamamoto et al., 2004; 2009; Menuet et al., 2007; Sifuentes-Romero et al., 2020). Increased Shh expression is in part responsible for the development of some regressive and constructive traits, namely eye degeneration (via lens apoptosis; Strickler et al, 2007), oral enhancement, and increased olfaction (Yamamoto et al., 2004; 2009; Pottin et al., 2011; Hinaux et al., 2016).

Cavefish have evolved in subterranean environments characterized by complete darkness, the absence of photosynthesis, reduced oxygen levels (hypoxia), and trophic challenges. Among these environmental factors, hypoxia has been proposed to play an important role in cavefish adaptative evolution (Boggs et al., 2021; 2022; van der Weele and Jeffery, 2022). The response to oxygen deficiency involves upregulation of Hypoxia Inducing Factor (HIF) genes, which encode transcription factors regulating hundreds of downstream target genes that are responsible for maintaining homeostasis according to oxygen availability (Semenza and Wang, 1992; Majmundar et al., 2010; Rashid et al., 2017; Corrado and Fontana, 2020). Accordingly, cavefish, but not surface fish, show constitutive overexpression of multiple *hif1a* genes and their downstream targets (van der Weele and Jeffery, 2022), indicating that they may have evolved a permanent canalized response to hypoxia. The *Astyanax* system provides a unique research platform in which the evolution of novel phenotypic traits can be studied from their beginning during surface fish colonization up to their current state in cavefish in the context of a relatively simple cave ecosystem (Poulson and White, 1969). In this study, we have used the *Astyanax* system to understand the evolution of enhanced hematopoiesis, a recently-discovered constructive trait in cavefish (van der Weele and Jeffery, 2022).

Hematopoiesis is the process in which erythrocytes and other blood cells are formed in two consecutive waves during vertebrate development. The first wave is known as primitive hematopoiesis, and the second wave is called definitive hematopoiesis (Keller et al., 1999; Davidson and Zon, 2004; Jagannathan-Bogdan and Zon, 2013). In teleosts, primitive hematopoiesis, which is the focus of the current study, begins with the formation of hematopoietic progenitor cells in the anterior and posterior lateral mesoderm during the tailbud stage (Paik and Zon, 2010; Gore et al., 2018). However, depending on teleost species, there are differences in the types of hemopoietic cells produced anteriorly and posteriorly. In zebrafish, the anterior lateral mesoderm (ALM) forms only macrophages, neutrophils, and microglia (Herbomel et al, 1999; Le Guyader et al., 2008), whereas the posterior lateral mesoderm is devoted to erythrocyte development. In contrast, the *Astyanax* ALM forms both macrophages and erythrocytes, and as in zebrafish the posterior lateral mesoderm produces only erythrocytes (van der Weele and Jeffery, 2022). It has been proposed that the expansion of erythrocyte development into the ALM may have been a pre-adaptation allowing *Astyanax* surface fish to colonize hypoxic subterranean habitats and evolve into cavefish (van der Weele and Jeffery, 2022). Cavefish show increased primitive hematopoiesis compared to surface fish: hematopoietic domains in both the ALM and posterior lateral mesoderm are increased in size and more larval erythrocytes are formed. Thus, primitive hematopoiesis is considered to be a constructive trait and an adaptive response to hypoxia in cavefish (van der Weele and Jeffery, 2022).

Here we explore the relationship between environmental hypoxia and Shh signaling in the evolution of primitive hematopoiesis and larval erythrocyte development using contemporary surface fish as a proxy for the ancestral cave colonizers. We show that hypoxia and increased Shh signaling enhance the size of primitive hematopoietic domains in the ALM and larval erythrocyte development in surface fish. Conversely, our results also indicate that inhibition of Shh signaling reduces the primitive hematopoietic domain in the ALM and decreases the number of erythrocytes in cavefish. Together, these results suggest a role for Shh signaling in *Astyanax* primitive hematopoiesis. Lastly, we demonstrate that hypoxia increases and expands midline Shh signaling itself in surface fish. These results provide evidence that the constructive evolution of cavefish primitive hematopoiesis was driven by a hypoxia-Shh axis.

## Materials and methods

### Biological materials

*Astyanax mexicanus* surface fish (Texas population) and cavefish (Pachón cave population) laboratory stocks were raised in a constant flow culture system. Embryos were obtained by natural spawning and reared at 23°C (Ma et al., 2021a). Fish husbandry protocols were approved by the University of Maryland Animal Care and Use Committee (IACUC #R-NOV-18–59) (Project 1241065–1) and conformed to National Institutes of Health guidelines.

### Hypoxia treatment

A hypoxia chamber (ProOx Model P110, BioSpherix, Parish, NY, USA) in which oxygen was reduced by nitrogen gas (HP, Airgas, Hyattsville, MD, USA) was used to expose embryos to a hypoxic environment. Embryos were exposed to 1 mg/L oxygen for about 6 hr beginning at 4 hours post-fertilization (50-60% epiboly). For hypoxia treatment embryos were placed in 12 or 24 well plates containing 2 or 3 ml of fish system water, respectively with 20-30 embryos per well (CytoOne, USA scientific, Ocala, FL, USA). Normoxic control embryos were raised in fish system-water outside of the hypoxia chamber. Hypoxia treatments lasted until embryos reached the tailbud stage (10 hpf), when some embryos were immediately prepared for RNA extraction or fixed for in situ hybridization and others were placed outside of the chamber in normoxic conditions and processed for RNA extraction or in situ hybridization after 4 hours. Some embryos were also kept until 36 hpf and then fixed for erythrocyte staining.

### SAG treatment

Shh signaling was increased by treating surface fish embryos with the cell permeable Smoothened agonist SAG (Cayman Chemical, Ann Arbor, MI, USA) (Chen et al., 2002a; Stanton and Peng, 2010). Stock solutions of SAG were prepared in DMSO. Embryos were dechorionated by treatment with 0.2 mg/ml protease type XIV (Sigma-Aldrich, St. Louis, MO, USA) for 1 min at room temperature, rinsed with fish system water and then treated with 0.5 μM SAG or DMSO (control) from 50% epiboly (4 hpf) to the tailbud stage (10 hpf). Embryos were then washed with and placed in fish system water until 14 hpf, when some embryos were processed immediately for RNA extraction or in situ hybridization and others were raised until 36 hpf for erythrocyte staining and quantification.

### Cyclopamine treatment

Shh signaling was inhibited by treatment of cavefish embryos with the Smoothened antagonist cyclopamine (Chen et al, 2002a). Cyclopamine stock solutions were prepared in 100% ethanol, and dechorionated embryos (see above) were treated with 30 μM cyclopamine (Cayman Chemical) or (Toronto Research Chemicals, Toronto, Canada) or ethanol (controls) using the same regime as described above for hypoxia and SAG treatments. At the end of the treatments (10 hpf), embryos were washed with and placed in fish system water, and fixed and processed for in situ hybridization at 14 hpf, or incubated until 36 hpf and processed for erythrocyte staining and quantification.

### In situ hybridization

In situ hybridization was performed on tailbud stage embryos fixed in 4% paraformaldehyde (PFA) and dehydrated in methanol using RNA probes for the *lmo2* and *shh* genes. *lmo2* was cloned from a 10 hpf cavefish cDNA library and *shh* from a 10 hpf surface fish cDNA library using the pCRII TOPO dual promoter vector (ThermoFisher Scientific, Waltham, MA, USA) transformed into One shot Top10 cells and the following primers: *lmo2* (ENSAMXG00000032986: 5’-ggcctctacaatcgagaggaaa-3’ and 5’ taccaagttgccgtttagtttgg-3’), and *shha* (ENSAMXG00000034481: 5’-cgaccgagacaagagcaaatac-3’ and 5’-aagttcccatccggtagagtag-3’). DIG-labeled probes were made using SP6 or T7 transcription kits (Roche, Mannheim, Germany) from linearized plasmid. In situ hybridizations were performed as described by Ma et al. (2014). Embryos were rehydrated stepwise into phosphate buffered saline (PBS), fixed with 4% PFA, digested with proteinase K (RPROTKSOL-R, Roche) post-fixed with 4% PFA, and hybridized at 60 °C for 16 hr. The hybridized specimens were washed with 2X and 0.2 X SSCT (150 mM sodium chloride; 15 mM sodium citrate; 0.1% Tween 2) followed by incubations in MABT blocking solution (Roche) and anti-DIG-AP Fab fragments (Roche). Specimens were rinsed in MABT buffer and PBS, equilibrated in AP buffer, and stained with BM-Purple (Roche). The stained specimens were imaged and photographed using a Zeiss Discovery V20 stereoscope with a Zeiss AxioCam HRc camera.

### RNA extraction and quantitative real time reverse transcriptase PCR

RNA was extracted with Trizol (ThermoFisher) from 30 embryos, treated with ezDNase, and cDNA was made with SuperScript IV Vilo Master mix (ThermoFisher) and used in quantitative real time reverse transcriptase PCR (qRT-PCR) with Takara SYBR Premix Ex Taq (Tli RNaseH Plus) (Takara Bio USA Inc, Mountain View, CA, USA) and LC480 (Roche) as described by van der Weele and Jeffery (2022). The mRNAs encoded by the *hypoxia-inducible factor* genes *hif1aa, hif1ab, hif1a-like*, and *hif1alike2* (equivalent to *hif3a*), the *sonic hedgehog* (*shha*) gene, and the downstream Shh-signaling system components *gli1* and *ptch1* were quantified using *gnb1b* as a reference. The primers used in qRT-PCR analysis are listed in Table 1. The ΔCt for each gene was calculated by subtracting the average Ct value of the reference gene. For graphical representation, the fold change was calculated as 2^-(ΔΔCt)^, where values > 1 show an increase and values < 1 a decrease in transcript levels. Variation was expressed as the range of fold change 2^-(ΔΔCt+stdevΔΔCt)^ for the upper value or 2^-(ΔΔCt-stdevΔΔCt)^ for the lowest value.

**Table 1.**
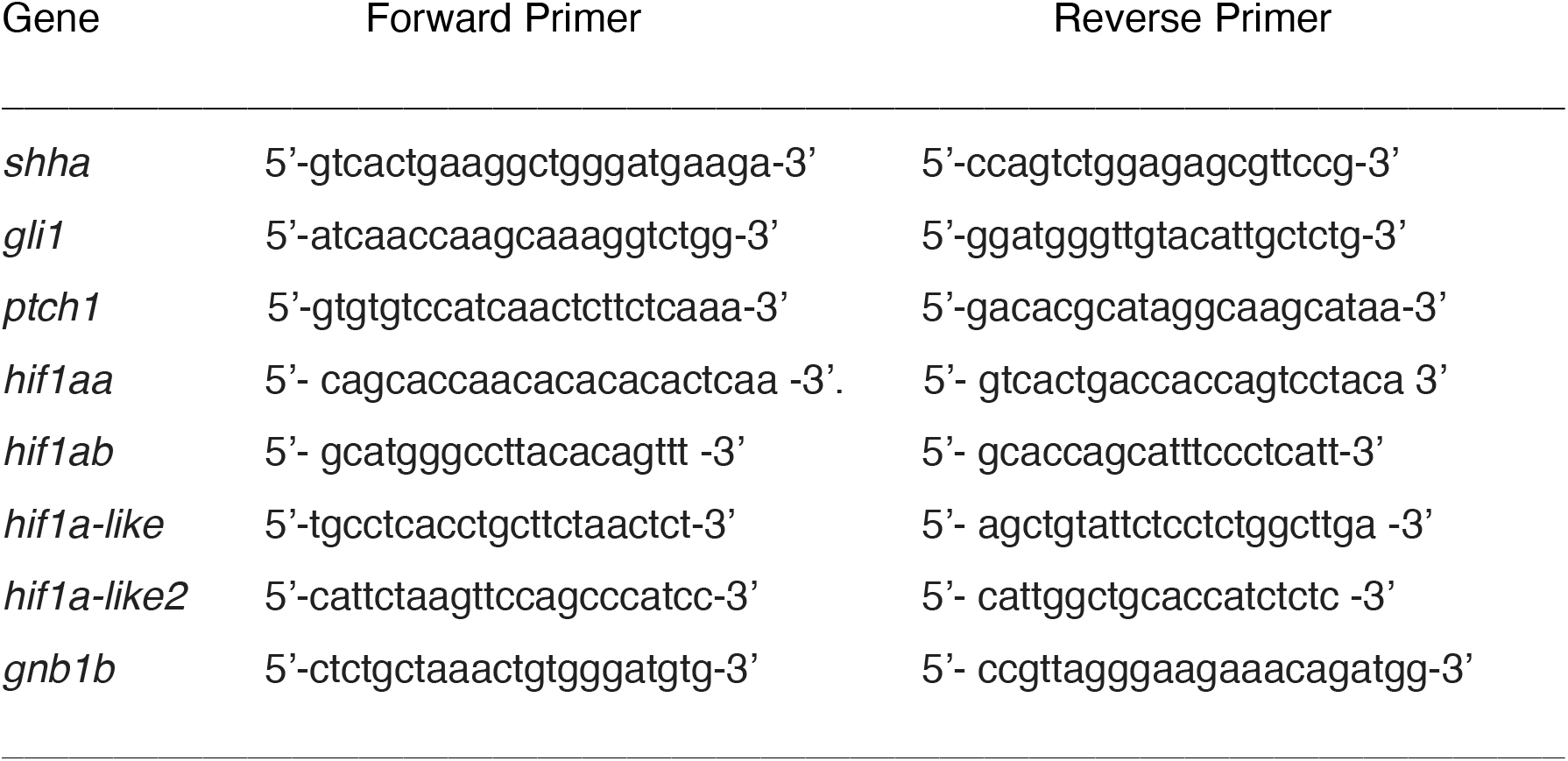
Oligonucleotide primer sequences used for qRT-PCR.

### Quantification of hematopoietic domains

The anterior hematopoietic domains were quantified as follows. For surface fish embryos, the number of *lmo2* stained cells was tabulated manually in the left and the right ALMs under a stereoscope and the counts were pooled for each embryo. For cavefish embryos, the widths of the *Imo2* stained ALMs were measured on the left and right side of the midline and then pooled for each embryo. Pooled values were used to account for differences in *lmo2* staining in the right and left ALMs in some embryos.

### Staining and quantification of erythrocytes

Erythrocytes were stained at 36 hpf with 0.6 mg ml^−1^ o-dianisidine (Sigma-Aldrich), 0.01 M sodium acetate (pH 4.5), 0.65% H_2_O_2_ for 15 min in the dark (Iuchi and Yamamoto, 1983). The stained embryos were rinsed in PBS, fixed in 4% PFA, and imaged. The number of stained blood cells were counted manually in an ROI (150 × 150 pixels) on the left and right Ducts of Cuvier and the totals were pooled for each larva. As described above, pooling of values was used to account for differences in erythrocyte number on the right and left sides of individual larvae.

### Quantification of Shh midline signaling domains

The width of Shh signaling domains were measured in the anterior and posterior midlines of surface fish from images using ImageJ-Fiji (Schindelin et al., 2019). Measurements of the width of the *shh* domain were made 50 pixels from either the anterior or posterior end of the domain.

### Statistics

Statistics were done using JMP pro 15 (SAS Institute Inc., Cary, NC, USA). Expression of the *hif1a* genes under normoxic or hypoxic conditions or *gli1* and *ptch1* in response to SAG treatment were compared using ΔCt in a three-way ANOVA (clutch x gene x treatment) followed by pairwise comparison of treatments for each gene using Student’s t-test (*hif1*-genes N = 6, *gli1* and *ptch1* N = 9, p<0.05). Expression of *shh* under normoxic and hypoxic conditions was analyzed using ΔCt with a two-way ANOVA (clutch x treatment) followed by Student’s t-test (N = 6, p<0.05).

Analysis of cell numbers in the anterior hematopoietic domain, the width of the hematopoietic domain, and the number of erythrocytes was done on pooled data from the left and right side of each fish using two-way ANOVA (clutch x treatment) followed by Student’s t-test where the lowest N was used to determine p-values (hematopoietic domain: N_normoxia_ = 41, N_hypoxia_ = 38, N_SAG_ = 14, N_DMSO_ = 18, N_ethanol_ = 100, N_cyclopamine_ = 98, erythrocyte number: N_normoxia_ = 80, N_hypoxia_ = 86, N_SAG_ = 48, N_DMSO_ = 48, N_ethano l_= 26, N_cyclopamine_ = 35, p<0.05).

The anterior and posterior widths of the *shh* domain were analyzed separately with two-way ANOVA followed by Student’s t-test where the lowest N was used to determine p-values (N_normoxia_ = 133, N_hypoxia_ = 98, p<0.05).

## Results

### Hypoxia increases primitive hematopoiesis and erythrocyte development in surface fish

The effects of hypoxia treatment on primitive hematopoiesis and erythrocyte development were determined in surface fish, which were used as a proxy for the cave colonizing ancestors of cavefish. Surface fish embryos were exposed to hypoxia from about 4 to 10 hours post-fertilization (hpf) and then returned to normoxic conditions (Fig. 1A). At 14 hpf RNA was extracted from embryos exposed to hypoxia (hypoxic embryos) and normoxic controls, and qRT-PCR was conducted to determine the effects on expression of the *hif1aa, hif1ab, hif1a-like*, and *hif1a-like2* genes, which encode transcription factors activated by oxygen depletion (Semenza and Wang, 1992; Majmundar et al., 2010) and are hallmarks of the hypoxia response in *Astyanax* (van der Weele and Jeffery, 2022). The results showed that the mRNA levels of all four *hif1a* genes were significantly increased in hypoxic embryos compared to normoxic controls (Fig. 1B), illustrating the effectiveness of the hypoxia treatment.

**Figure 1.**
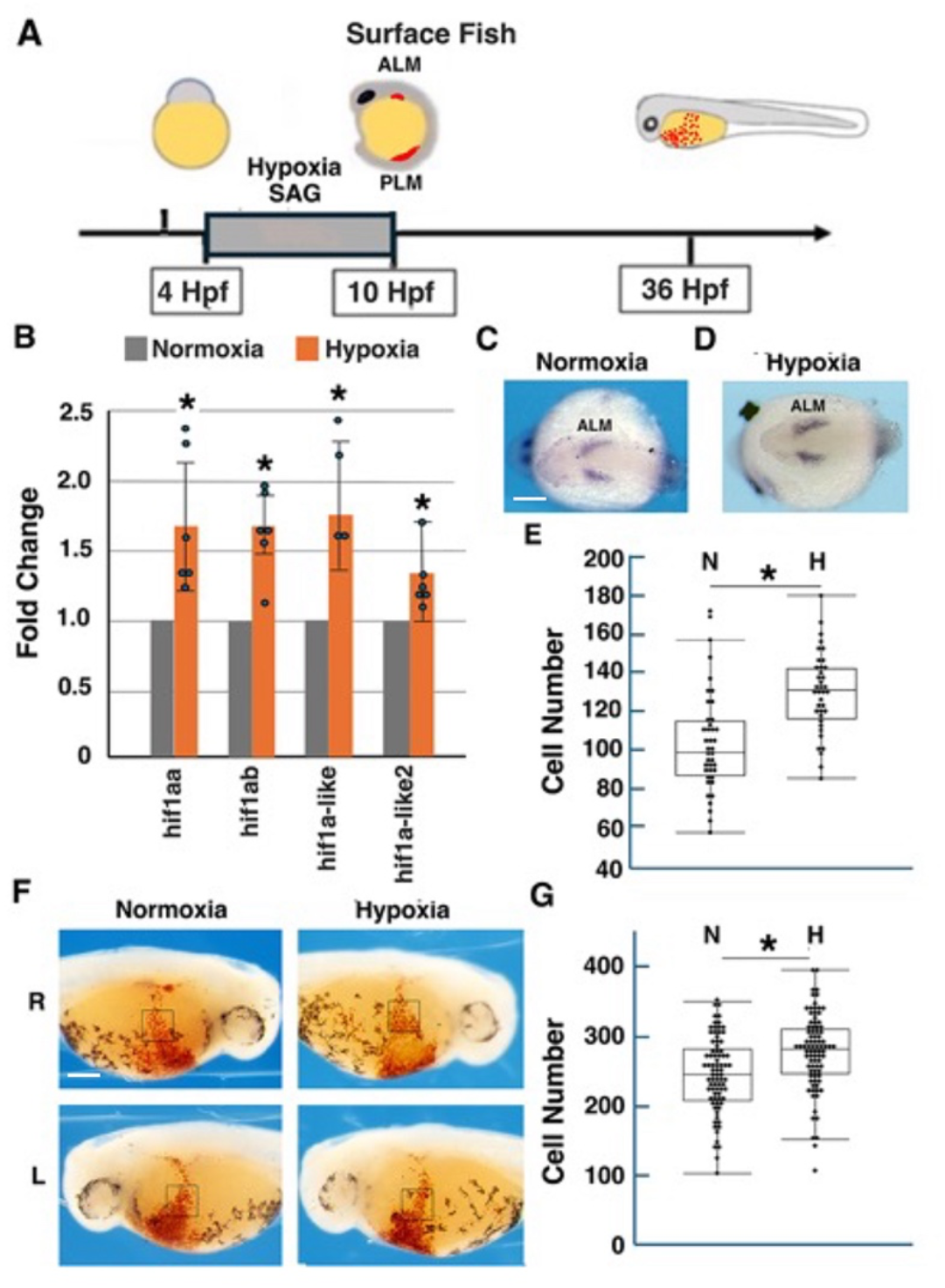
Hypoxia increases primitive hematopoiesis and erythrocyte development in surface fish. A. The time course of hypoxia (B-G) and SAG (Fig. 2), treatment in surface fish showing hematopoietic domains in the anterior lateral mesoderm (ALM) and posterior lateral mesoderm (PLM). Hpf: Hours post-fertilization. B. Quantification of *hif1a* gene expression by qRT-PCR in hypoxic relative to normoxic surface fish at 14 hours post-fertilization (hpf). Bars show the mean and standard errors of fold changes in *hif1aa, hif1ab, hif1-a-like*, and *hif1a-like2* mRNA levels. Asterisks from left to right: p =.006, p = .036, p = .009, and p = .049. C-E. Effects of hypoxia on primitive hematopoiesis in surface fish. C, D. In situ hybridization showing *Imo2* staining of the bilateral ALM domains of hypoxic embryos (D) and normoxic controls (C) at 14 hpf. Anterior poles are on the left. Scale bar: 200 μm; magnification is the same in both frames. E. Bar graphs showing the mean numbers and standard errors of *lmo2* stained cells in the bilateral ALMs of normoxic (N) and hypoxic (H) embryos at 14 hpf. Asterisk: p < .0001. F. o-dianisidine staining of erythrocytes in the Duct of Cuvier at 36 hpf on the left (L) and right (R) sides of larvae exposed to hypoxia (H) during embryogenesis (see A) and normoxic (N) controls. Scale bar: 200 μm; magnification is the same in all frames. Boxes: erythrocyte counting areas. G. Bar graphs showing the mean numbers and standard errors of o-dianisidine stained erythrocytes of larvae exposed to hypoxia during embryogenesis (H) and normoxic controls (N) at 36 hpf. Asterisk: p < .0001.

To determine the effects of hypoxia on hematopoiesis, we used in situ hybridization with the *LIM-only 2* (*lmo2*) gene, which encodes a transcription factor required for erythrocyte development (Patterson et al., 2007; Gering et al., 2003) and strongly labels *Astyanax* hematopoietic domains (van der Weele and Jeffery, 2022). As shown in Figure 1C, D, *lmo2* staining of 14 hpf embryos revealed bilateral hematopoietic domains, the sites of erythrocyte formation in the anterior lateral mesoderm (ALM) of surface fish and cavefish embryos (van der Weele and Jeffery, 2022). Quantification showed that the mean number of *Imo2*-stained cells in the ALM was significantly increased in hypoxic relative to normoxic embryos (Fig. 1E), indicating that hypoxia treatment enhances hematopoiesis in surface fish.

We next asked if hypoxia treatment affects larval erythrocyte development. Surface fish were exposed to hypoxia (Fig. 1A), hypoxic embryos and normoxic controls were raised until 36 hpf, when the heart begins to beat and the first blood cells stream over the yolk mass through the Duct of Cuvier (van der Weele and Jeffery, 2022), then the larvae were fixed and stained with the erythrocyte-specific marker o-dianisidine (Iuchi and Yamamoto, 1983). Quantification of o-dianisidine-stained cells indicated a significant increase in hypoxic compared to normoxic surface fish (Fig. 1G). The results demonstrate that exposure to hypoxia increases primitive hematopoiesis and erythrocyte development in surface fish.

### Sonic Hedgehog upregulation increases primitive hematopoiesis and erythrocyte development in surface fish

Shh signaling is expanded along the embryonic midline in cavefish compared to surface fish (Yamamoto et al., 2004; 2009; Menuet et al., 2007) and is in a position to affect hematopoiesis in the nearby bilateral hematopoietic domains. Therefore, we next asked whether increased Shh signaling affects primitive hematopoiesis and larval erythrocyte numbers in surface fish. We used a pharmacological approach via the Smoothened agonist SAG (Chen et al., 2002a, b) to upregulate Shh signaling. SAG treatment began at 4 hpf and was continued until tailbud stage (10 hpf) (Fig. 1A), then the treated embryos were rinsed several times into fresh fish system water and allowed to develop to 14 or 36 hpf. Controls were subjected to the same treatment in DMSO, the vehicle for SAG solution. To confirm the effectiveness of SAG treatment in upregulating the Shh signaling system, we used qRT-PCR to determine the expression of the downstream *gli1* and *ptch1* genes. The results showed that expression of both genes was significantly increased in SAG treated compared to control embryos (Fig. 2A), confirming Shh pathway upregulation.

**Figure 2.**
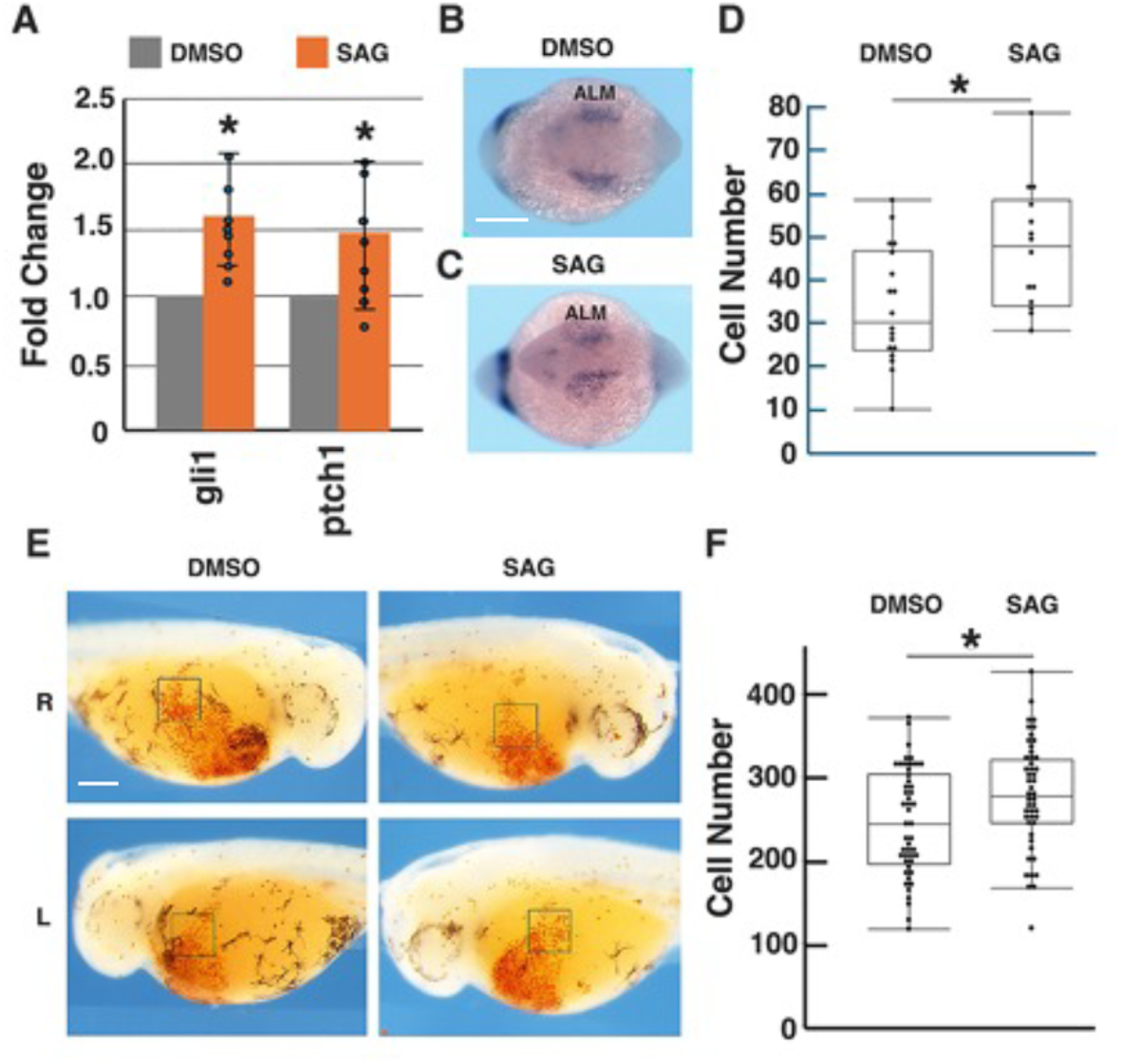
Sonic Hedgehog upregulation increases primitive hematopoiesis and erythrocyte development in surface fish. A. RT-qPCR showing the effects of SAG on the downstream Shh signaling genes *gli1* and *ptch1* in tailbud embryos. Asterisks from left to right: p = 003; p= .046. B-D. Effects of SAG on the number of *Imo2* stained cells in the anterior lateral mesoderm (ALM). B, C. In situ hybridization showing *Imo2* staining of the ALM in SAG treated and DMSO control tailbud embryos. Scale bar: 200 μm; magnification is the same in both frames. D. Bar graphs comparing the number of *lmo2* stained cells in the anterior hematopoietic domains of SAG treated and control embryos. Bars represent the mean numbers of stained cells in the left and right ALM with standard errors. N_SAG_ = 14. N_DMSO_ = 18. Asterisk: p = .0069. E. o-dianisidine staining of erythrocytes in the Duct of Cuvier at 36 hpf on the left (L) and right (R) sides of larvae exposed to SAG during embryogenesis (see Fig. 1A) and controls. Boxes: erythrocyte counting areas. Scale bar: 200 μm; magnification is the same in all frames. F. Bar graphs showing the mean numbers and standard errors of o-dianisidine stained erythrocytes in the pooled right and left sides of 36 hpf larvae that developed from SAG treated embryos and ethanol treated controls (see Fig. 1A). Asterisk: p = .0005.

The effects of Shh upregulation on *Imo2* staining in the ALM at 14 hpf and erythrocyte development at 36 hpf are shown in Fig 2B-F. SAG treatment significantly increased the numbers of *lmo2* expressing cells in the ALM (Fig. 2B-D) and o-dianisidine labeled erythrocytes later in development (Fig. 2E, F). The results indicate that upregulation of the Shh pathway by SAG increases primitive hematopoiesis and larval erythrocyte development in surface fish.

### Sonic hedgehog inhibition decreases primitive hematopoiesis and erythrocyte development in cavefish

Shh midline signaling, primitive hematopoiesis, and erythrocyte development is enhanced in cavefish relative to surface fish (Yamamoto et al, 2004; 2009; Menuet et al., 2007; Sifuentes-Romero et al., 2020; van der Weele and Jeffery, 2022). To determine whether Shh is involved in this enhancement, Shh signaling was inhibited with the Smoothened antagonist cyclopamine (Fig. 3A). The effects of cyclopamine on *Imo2* staining in the ALM and erythrocyte development are shown in Figure 3H-L. In contrast to surface fish, the strong enhancement of *Imo2* staining in the cavefish ALM, did not permit the resolution and quantification of individual cells. Thus, *lmo2* staining was compared in cyclopamine-treated and control cavefish embryos by measuring the lateral widths of ALMs. Inhibition of the Shh pathway significantly decreased the mean width of *Imo2* stained ALMs in 14 hpf cavefish embryos (Fig. 3C, D) and also significantly reduced the numbers of o-dianisidine labeled erythrocytes in 36 hpf cavefish larvae (Fig. 3E-F). These results indicate that downregulation of Shh signaling by cyclopamine decreases the ALM hematopoietic domain and erythrocyte numbers in cavefish.

**Figure 3.**
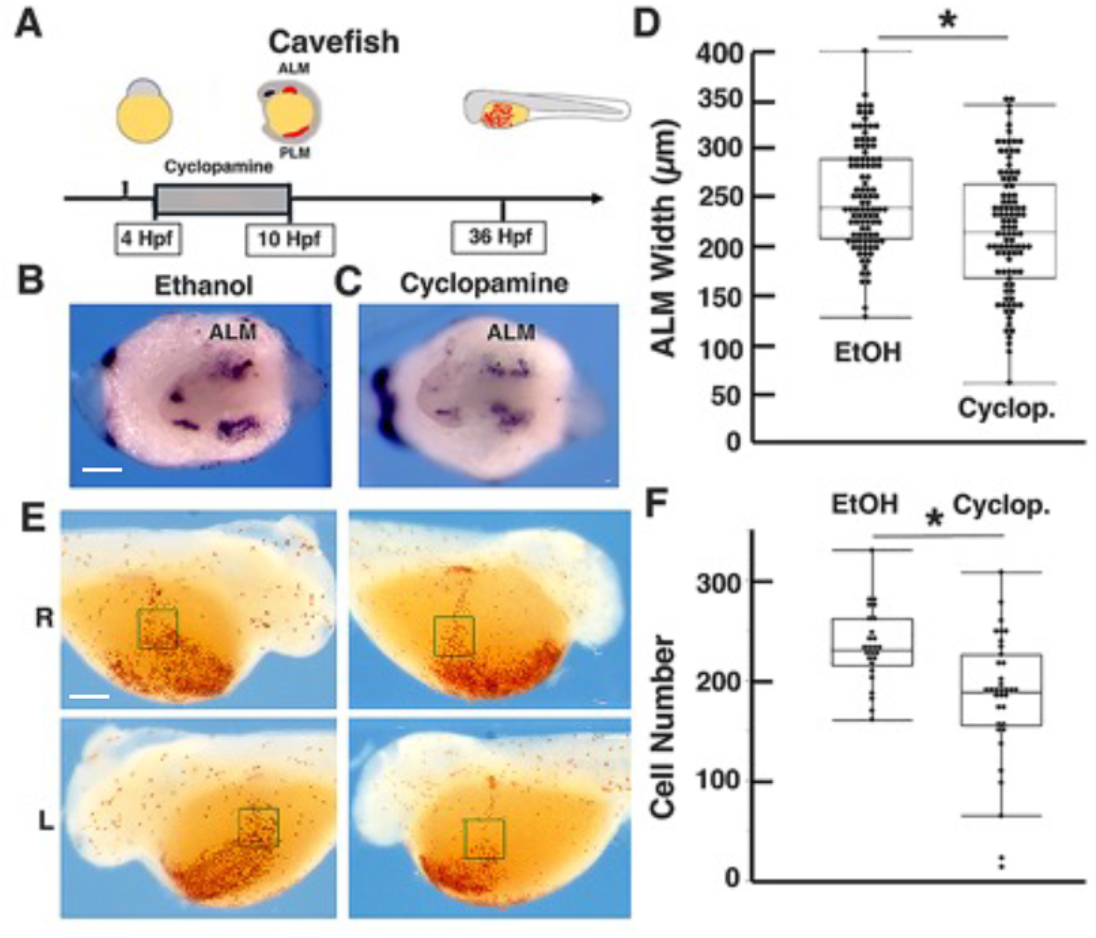
Sonic Hedgehog inhibition decreases primitive hematopoiesis and erythrocyte development in cavefish. A. The time course of cyclopamine treatment in cavefish showing enlarged hematopoietic domains in the anterior lateral mesoderm (ALM) and posterior lateral mesoderm (PLM). B-D. Effects of cyclopamine on the width of the *Imo2* stained ALM in tailbud embryos. B, C. In situ hybridization showing *Imo2* staining in the ALM of cyclopamine-treated tailbud embryos (C) and ethanol (EtOH)-treated controls (B). Scale bar: 200 μm; magnification is the same in all frames. D. Bar graphs comparing the width of *lmo2* stained anterior hematopoietic domains in cyclopamine and ethanol treated embryos. Bars represent the pooled means of left and right ALM widths with standard errors. N_ethanol_ = 100. N_cyclopamine_ = 98. Asterisk: p < .0001. E. o-dianisidine staining of erythrocytes in the Duct of Cuvier at 36 hpf on the left (L) and right (R) sides of larvae exposed to cyclopamine during embryogenesis (see A) and ethanol treated controls. Boxes: erythrocyte counting areas. Scale bar: 200 μm; magnification is the same in all frames. F. Bar graphs showing the mean numbers and standard errors of o-dianisidine stained erythrocytes in the pooled right and left sides of larvae exposed to cyclopamine during embryogenesis (see A) and controls at 36 hpf. Asterisk: p < .001.

Taken together, the SAG (Fig. 2) and cyclopamine (Fig. 3) results suggest that primitive hematopoiesis and erythrocyte development is controlled by Shh signaling in *Astyanax* and that increased Shh signaling enhances both of these traits in cavefish.

### Hypoxia increases Sonic Hedgehog signaling along the surface fish embryonic midline

The results described above opened the possibility that hypoxia treatment may affect primitive hematopoiesis and erythrocyte development by enhancing the Shh signaling pathway. To test this hypothesis, surface fish were exposed to hypoxia using the regime depicted in Figure 1A and the effects on *shh* mRNA expression were compared by qRT-PCR and in situ hybridization of hypoxic tailbud embryos and normoxic controls. Hypoxic embryos showed a significant increase in *shh* mRNA levels compared to normoxic counterparts (Fig. 4A).

**Figure 4.**
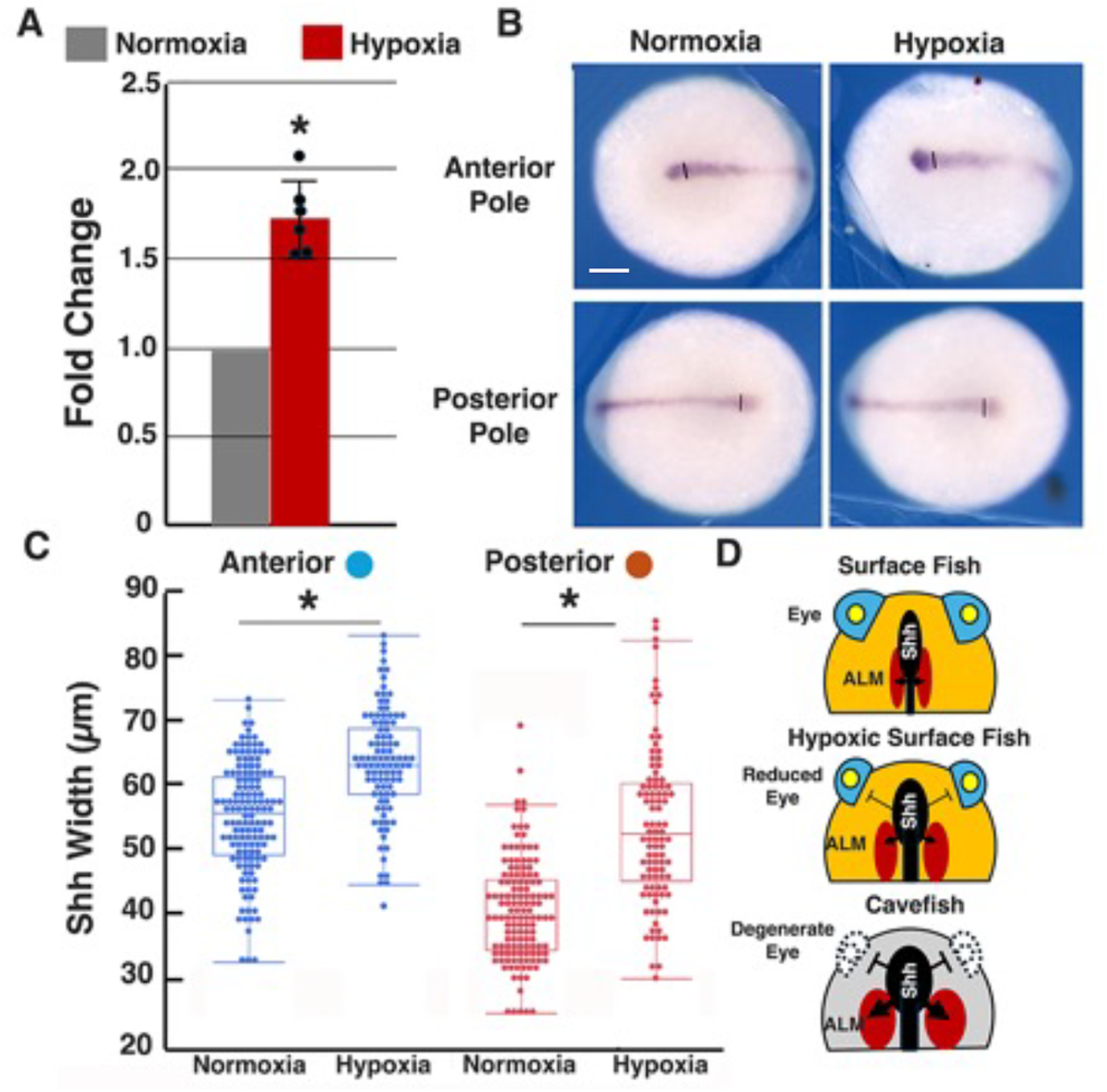
Hypoxia increases Sonic Hedgehog signaling in surface fish embryos. Quantification of *shh* gene expression by qRT-PCR in hypoxic relative to normoxic surface fish embryos at 10 hours post-fertilization (hpf). Bars show mean and standard errors of fold changes *shh* mRNA levels. Asterisk: p = .0001. B. In situ hybridization showing *shh* expression along the midline of normoxic and hypoxic surface fish embryos at the tailbud stage. Vertical lines: positions along the midline where expression domain width was measured. Scale bar: 200 μm; magnification is the same in all frames C. Quantification of *shh* expression-domain width differences in the anterior and posterior midlines of hypoxic and normoxic embryos. Asterisks: p < .0001. D. A model for the effects of hypoxia on midline *shh* signaling and the size of anterior hematopoietic domains in surface fish and cavefish. According to the model, hypoxia negatively affects eye development (van der Weele and Jeffery, 2022) and positively affects primitive hematopoiesis (present investigation) in surface fish and cavefish though its effects on Shh midline signaling.

Furthermore, the width of the *shh* expression domain was significantly increased in both anterior to posterior locations along the midline of tailbud embryos (Fig. 4B, C). These results, along with the results obtained above, suggest that hypoxia increases primitive hematopoiesis and erythrocyte development by enhancement of the Shh midline signaling system.

## Discussion

The subterranean habitats of *Astyanax* cavefish are characterized by complete darkness, lack of photosynthesis, and hypoxia (Rohner et al., 2013; Ornelas-Garcia et al., 2018). Unusual phenotypes, such as the loss of eyes, depigmentation, obesity, and novel behaviors (Jeffery, 2020; Kowalko, 2020; Perera et al. 2023), evolved under these conditions, which may have allowed cavefish to adapt and survive in challenging environments. As an adaptation to reduced oxygen, erythrocyte development may have been enhanced in cavefish compared to surface fish (Boggs et al., 2022; van der Weele and Jeffery, 2022), a trait shared with animals adapted to living at high altitudes (Acton and Harvey, 1912; Haase, 2013). The present study aimed to understand the role of reduced oxygen in the evolution of cavefish hematopoiesis using contemporary *s*urface fish as a proxy for the original cave colonizers. Accordingly, surface fish embryos were exposed to hypoxia in the laboratory and the effects on the first wave of hematopoiesis and subsequent development of larval erythrocytes were determined. Our results show that hypoxia mimics the cavefish hematopoietic phenotype by promoting the formation of larger hematopoietic domains expressing the *Imo2* marker gene and the development of more erythrocytes in surface fish. We further show that primitive hematopoiesis is controlled by the Shh pathway, a signaling system that is also responsible for the enhancement of other cavefish traits (Yamamoto et al., 2004; 2009; Pottin et al., 2011; Hinaux et al., 2016; Sifuentes-Romero et al., 2020). In addition, the Shh system itself can be elevated by exposure of surface fish embryos to hypoxia in the laboratory. These results suggest that the evolutionary enhancement of primitive hematopoiesis in cavefish is controlled by a hypoxia-Shh axis.

Classic studies indicate that the increases in the production of hemoglobin, the ability of hemoglobin to bind oxygen, and/or the development of more erythrocytes are adaptations to reduced oxygen levels in animals living at high altitudes (Windsor and Rodway, 2007; Storz, 2007; Webb et al., 2022). However, the early developmental mechanisms underlying these hypoxia dependent changes are not well known. Comparative studies in *Astyanax* have shown that bilateral hematopoietic domains representing the earliest stages of the hematopoietic first wave are larger and contain more *Imo2* expressing cells in cavefish than in surface fish (van der Weele and Jeffery, 2022). Furthermore, as known for other cavefish traits (Ma et al, 2018; 2021b; Torres-Paz et al., 2019), the discovery that cavefish erythrocyte enhancement is a maternal effect extends the developmental origin of hematopoietic changes into oogenesis (van der Weele and Jeffery, 2022). The present study demonstrates that early events in hematopoietic development can be modified by reduced oxygen in the surface fish proxy for cave colonizers, suggesting that environmental hypoxia is a driver for hematopoietic evolution. Thus, evolutionary changes driven by low oxygen levels in subterranean environments (Malard and Hervant, 2001) may be analogous to adaptations in surface habitats characterized by high altitude or other hypoxic habitats.

Our results provide evidence suggesting that the Shh signaling system, which is known to be involved in the regulation of cavefish traits such as gain in olfaction and feeding structures and loss of eyes (Yamamoto et al., 2004; 2009; Menuet et al., 2007; Pottin et al., 2011; Hinaux et al., 2016), has additional roles in the enhancement of primitive hematopoiesis and erythrocyte development. Accordingly, upregulating Shh signaling in surface fish by SAG treatment increased primitive hematopoiesis, as shown by larger hematopoietic domains and the development of more erythrocytes, whereas inhibiting Shh signaling in cavefish by cyclopamine treatment decreased primitive hematopoiesis, as shown by smaller hematopoietic domains and fewer larval erythrocytes. Conflicting information has been published concerning the role of Hedgehog signaling in primitive and definitive hematopoiesis in vertebrates, and research with different species have provided opposing results (Bhardwaj et al., 2001; Hoffman et al., 2009; Lim and Matsui, 2010; Chen et al., 2023). Although most previous studies support the role of Shh signaling in definitive hematopoiesis (Gering and Patient, 2005), the function of Shh signaling in primitive hematopoiesis is poorly understood. Thus, our studies in *Astyanax* embryos have filled a gap in understanding of the developmental regulation of blood development by demonstrating a role for Shh signaling in the first wave of hematopoiesis.

The primary purpose for conducting the present study was to further understand the control of evolutionary changes in primitive hematopoiesis and erythrocyte development in cavefish. An important breakthrough was the finding that hypoxia, in addition to enhancing the first wave of erythrocyte development, also increases Shh signaling along the embryonic midline. Accordingly, surface fish embryos exposed to hypoxia showed a significant increase in *shh* mRNA levels and lateral expansion of *shh* expression, thus mimicking Shh overexpression in cavefish (Yamamoto et al., 2004; 2009, Menuet et al., 2007; Sifuentes-Romero et al., 2020) and revealing a potentially important link between hypoxia and cavefish traits that are sensitive to low oxygen levels, including primitive hematopoiesis. To our knowledge this is the first report of an effect of hypoxia on the Shh signaling system during vertebrate embryogenesis. However, the effects of hypoxia and the HIF pathway on Shh signaling in adult tissues, organs, and tumors has been documented previously (Hwang et al., 2008; Bijlsma et al., 2009; Wang et al., 2010; Spivak-Kroizman et al., 2013; Katagiri et al., 2018; Bhuria et al., 2019) and supports our results in *Astyanax* embryos. Thus, we propose that a hypoxia-Shh axis may have been the driving force of hematopoietic evolution in cavefish. A key factor in the hypoxia-Shh axis is likely to be activation of the HIF regulatory system by oxygen deficiency, and this assumption is supported by up-regulation of four different *hif1a* genes in surface fish embryos exposed to hypoxia and constitutive upregulation of some of these genes in cavefish (Fig. 1B and van der Weele and Jeffery, 2022). How the HIF system may influence Shh signaling is presently unknown: it is possible that the Shh pathway is either a direct or indirect target of HIF1 transcription factors. Whichever the case, it will be important in future studies to focus on the role of the HIF1 regulatory system in the evolution of cavefish hematopoiesis and Shh-driven cavefish traits.

Previous studies suggested that the evolution of cavefish eye degeneration, a regressive trait caused in part by Shh induction of lens apoptosis (Strickler et al., 2007), may be linked in a pleiotropic relationship with Shh-dependent constructive traits, such as oral size and tastebud enhancement (Yamamoto et al., 2009). Accordingly, increased Shh signaling along the anterior midline could simultaneously enhance oral traits and reduce eye development, thus presenting a potential target for natural selection. The present results call for a revised version of the pleiotropic hypothesis to include the evolution of the cavefish hematopoietic system, which develops bilaterally in the ALM flanking the *shh* expressing midline and could be another pleiotropic target for natural selection. The revised hypothesis featuring the positive effects of increased Shh signaling on hematopoiesis coupled with the negative effects on eye development is illustrated in Figure 4D. Previous studies have also demonstrated a role for hypoxia in *Astyanax* eye degeneration (Ma et al., 2020; van der Weele, 2022), which is in line with the pleiotropic model.

In conclusion, the current study has increased our understanding of how a subterranean environment characterized by oxygen deficiency could drive cavefish evolution. Our research has revealed key roles of hypoxia and Shh signaling in the development and evolution of primitive hematopoiesis and larval erythrocyte enhancement in the *Astyanax* model, discovered the role of a novel hypoxia-Shh axis on enhanced hematopoiesis, and opened the possibility of primitive hematopoiesis as a trait that may be relevant to the evolution of cavefish eye degeneration by indirect selection.

## Acknowledgements

We thank Ruby Dessiatoun and Karina Lacroix for maintenance of the fish facility and Mandy Ng for technical assistance

## Author contributions

CMW and WRJ conceived the project, CMW, KCH, and KCR performed the research, WRJ and CMW wrote the manuscript with input from the co-authors.

